# Antibiotic tolerance due to filamentation shapes β-lactam pharmacodynamics in *Escherichia coli*

**DOI:** 10.64898/2026.08.25.747073

**Authors:** Aryan Ramachandran, Josia Pool, Arjan de Visser, Hilje Doekes, Aditi Batra

## Abstract

Pharmacodynamic curves describe how changes in drug concentration affect pathogen growth. They are essential for designing treatments that promote pathogen eradication and minimize the evolution of antibiotic resistance. The classical function for modelling pharmacodynamics is a phenomenological, S-shaped curve with stable growth and death rates separated by a single drop. In this study, we characterized the pharmacodynamic curve of the β-lactam antibiotic cefotaxime (CTX) acting against *Escherichia coli*. We found that the relationship between CTX concentration and net growth rate diverged from classical model predictions, instead yielding a two-step curve defined by distinct phases of growth, population maintenance, and killing. We hypothesized that the intermediate phase arose from antibiotic tolerance conferred by bacterial filaments. Microscopic assessment of treated cells indeed showed a difference in degree of filamentation with concentration. We further sought to explain this with a semi-mechanistic pharmacodynamic function, modelling the binding of CTX to its cellular targets, penicillin binding proteins (PBP) 1 and 3. By incorporating the preferential concentration-dependent binding of CTX to PBP3 and then PBP1, yielding filaments or lysed cells respectively, we replicated the two-step curve in silico. We also assessed the pharmacodynamics of CTX against mutants conferring resistance; these displayed further altered curves, in line with their fitness costs. Altogether, our results show that CTX has a two-step pharmacodynamic curve against *E. coli* arising from multiple targets separated in their affinity for the antibiotic. We present a model offering a mechanistically grounded framework for capturing such dynamics. These pharmacodynamic curves deserve careful consideration when defining optimal dosing.

**Importance:** β-lactams make up the most prescribed group of antibiotics in the world. As infecting bacteria evolve resistance, however, these antibiotics become less effective in treating infections. Designing rational treatments to optimise therapy is therefore paramount. In our research, we investigate the effects of increasing antibiotic concentrations on bacterial survival, an important aspect of designing treatment plans. We study the action of the β-lactam antibiotic cefotaxime against the bacterium *Escherichia coli*. By using a combination of laboratory experiments and computer simulations, we demonstrate an unusual relationship between antibiotic concentration and bacterial growth stemming from antibiotic tolerance provided by filamentous cells. These findings improve our understanding of how bacteria respond to β-lactam antibiotics, and highlight the importance of accounting for such effects when designing treatments to fight resistant infections.

## Introduction

The evolution of antimicrobial resistance (AMR) poses a threat to global health. As microbial populations undergo antibiotic exposure, resistant variants may evolve and spread, resulting in treatment failure. Such failure can be severe: bacterial AMR directly caused 1.27 million deaths globally in 2019 (Murray et al., 2022). Rational treatment design to counter resistance is imperative.

An important factor to consider is how antibiotic dosage affects the net rate of change of the bacterial population, also known as pharmacodynamics. The popular pharmacodynamic model is that of Regoes et al., published in 2004, which still finds use more than 20 years later(Andersson et al., 2026; Ankomah & Levin, 2014; Childers et al., 2026; Day & Read, 2016; Regoes et al., 2004). This model is phenomenological, describing a relationship where the antibiotic has little effect on bacterial numbers at low concentrations, followed by a steep reduction in growth rates at the MIC and an eventual stabilization of a maximum killing rate at high concentrations. Its use now extends beyond classical dosing regimens: it has been used to define mutant selection windows (Witzany et al., 2023), and has been built on to develop models exploring successful combination treatments (Nyhoegen et al., 2024).

β-lactam antibiotics, are the most prescribed class of antibiotics in the world; accounting for more than half of all antibiotics in use (Elander, 2003; Sargianou et al., 2025). Their popularity comes from their high efficacy and excellent safety profile (MacVane et al., 2014). β-lactams act by binding to Penicillin Binding Proteins (PBPs), a set of enzymes integral to peptidoglycan synthesis and cell division in bacteria. Binding of the antibiotic inactivates the PBPs ultimately resulting in cell death (Hayes & Orr, 1983; Tuomanen et al., 1986). Amongst the β-lactams, the third-generation cephalosporin cefotaxime (CTX) stands out due to its broad-spectrum activity and stability against a wide range of β-lactamases (Todd & Brogden, 1990). CTX exerts its bactericidal effects through preferential binding to distinct PBPs at different concentrations. At lower concentrations, CTX primarily targets PBP3, the protein responsible for septal formation between dividing daughter cells (Nguyen-Distèche et al., 1998). Binding of CTX to PBP3 inhibits cell division while growth continues, resulting in the formation of long cylindrical filaments. Prolonged filamentation ultimately results in loss of structural viability and non-lytic cell death (Hayes & Orr, 1983; Kim et al., 2023; Tuomanen et al., 1986). At higher concentrations, CTX additionally binds to PBP1a and PBP1b, proteins essential for cell wall integrity. Their inactivation interferes with cell wall regeneration and causes lytic death (Pazos & Vollmer, 2021; Yousif et al., 1985).

Earlier works describing the pharmacodynamics of CTX have focused on its effects over extended periods of time, studying only a limited range of concentrations (Alexandersen et al., 2025; Frimodt-MØLler et al., 1987; Visalli et al., 1996). In this study, we aimed to characterize the time-killing dynamics of CTX in *E. coli* REL606 and its mutants with defined resistance mechanisms, assessing pharmacodynamics over a wider range of concentrations. We find that the net rate of population turnover, as a function of antibiotic concentration, follows two-step kinetics. A semi-mechanistic computational model incorporating multiple targets accounts for the observed kinetics.

## Methods

### Study Systems

Four strains of *E. coli* that are isogenic except for the presence and expression of three different cefotaxime (CTX) resistance mechanisms were used (Schenk et al., 2022). The CTX-susceptible *E. coli* B REL606-CFP served as the wild type (Dillon et al., 2016). The import mutant *ΔompF*, has the gene encoding the outer membrane porin OmpF deleted. The primary channel for the influx of CTX (Jaffe et al., 1982), OmpF’s deletion reduces the rate at which the antibiotic enters the cell, conferring resistance. The export mutant, *ΔacrR*, has *acrR*, a gene coding for the negative regulator for the AcrAB-TolC multidrug efflux system deleted, resulting in overexpression of the pump and resistance through expulsion of CTX from the cell (Nagano & Nikaido, 2009; Weston et al., 2018). The final mutant, *“*G238S*”*, expresses high levels of a mutated TEM-1 β-lactamase from a multicopy plasmid, pACTEM. The mutated version of the β-lactamase, has higher activity against CTX than the native version, conferring resistance against the antibiotic. The expression of the gene is induced by isopropyl-β-D-thiogalactopyranoside (IPTG) and the plasmid maintained through selection with tetracycline (Salverda et al., 2017). These mutants were chosen because they were observed to be the dominant resistance mechanisms that evolved in *E. coli* REL606 under CTX stress (Schenk et al., 2022).

### Time-kill Assays

Time-kill assays provide an understanding of bacterial growth and death rates under antibiotic stress (Alexandersen et al., 2025; Ferro et al., 2015; Foerster et al., 2015). Bacteria were treated with a range of CTX concentrations and the change in viable cell number was assessed over 3 hours. A single run included 16 antibiotic concentrations: one growth control with no CTX, and 15 concentration doublings. The maximal concentration assessed for the WT, *ompF*, and *acrR* lines was 250 µg/mL. Owing to the markedly increased resistance of the G238S strain, a concentration range up to 5000 µg/mL was used. Additionally, to ensure the carriage and expression of the plasmid in the G238S strain, the cultures also contained 10 µg/mL tetracycline and 50 µM IPTG. Two independent runs of the assay were performed per strain. Every run, in turn included three biological replicates. An inoculum of 5 × 10^7^ CFU/mL was used for the assays, leaving sufficient room for both growth and death to be measured. For each concentration, the required volume from the overnight cultures was pelleted and then resuspended in 500 µl M9 medium (6.78 g/L Na_2_HPO_4_.2H_2_O, 3 g/L KH_2_PO_4_, 1 g/L NH_4_Cl, 0.5 g/L NaCl, 0.12 g/L MgSO_4_, 0.015 g/L CaCl_2_, 4 g/L D-Glucose-Monohydrate, 2 g/L Cas-amino acids) containing CTX at the desired concentrations. 100 µL of these cultures were inoculated into the wells of 96 well plates (growth plates) and incubated at 37°C and 220 rpm. Every hour (including the initial 0-hour timepoint), a well from the growth plates was sacrificed, transferred to a different 96 well plate) and diluted in seven 10-fold steps. Following serial dilution, 7 µl from each well was spot plated onto 120 mm square M9 agar plates. The plates were incubated at 37 °C overnight and the colonies counted the following day. Only spots with greater than 4 and less than 50 colonies were counted. Change in colony forming units (CFU) over time was extrapolated directly from CFU counts for each timepoints. The growth/death rates for each concentration were obtained by fitting a linear regression through the natural log of the CFU counts versus time. CFU counts less than 1400 per mL were excluded as they lay below the assay’s limit of detection.

### Microscopic Imaging

To probe the mechanism behind our observations from the time-kill experiments, cell morphology was studied using microscopy. Each microscopy assay involved culturing cells in M9 medium with different CTX concentrations, as in the time-kill assays. 6 concentrations were assessed per assay, with 3 biological replicates. The concentrations were chosen to represent the different phases observed in the pharmacodynamic curve. At the time of inoculation (Timepoint 0), only the growth control was imaged as we assumed no significant changes occurred in such a short time span. Cells from all concentrations were then imaged at 45 (Timepoint 1) and 90 (Timepoint 2) minutes post inoculation. To facilitate faster imaging while retaining morphological information from independent replicates, all 3 biological replicates were pooled together. The pooled samples were spun down and all but 20 µl of the supernatant discarded. After resuspension, 10 µl of this was pipetted onto a glass slide and a clean coverslip was placed on top. We thus had one slide for each concentration assessed at every timepoint. The cells were imaged using a Leica DFC340 FX microscope using the F36-544 CFP channel. Using the 40x objective lens, 6 distinct areas were imaged for each concentration and timepoint.

### Image Analysis

The images were then analysed to categorise cell morphologies and measure their frequencies. The imaged cells were converted into measurable entities or “masks” (cell boundaries marked) using the cellular segmentation algorithm Cellpose (Stringer et al., 2021). The masks were then processed via Fiji (Schindelin et al., 2012), correcting any improperly segmented cells. The Area, Perimeter, and Circularity of each cell was measured using pixels as units. Area is the total number of pixels housed within a mask while Perimeter measures the pixels comprising the mask (or cell) boundary. Circularity assesses how close to a true circle the cell shape is; it is calculated using the measured values for Area (*A*) and Perimeter (*P*) through the formula 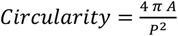. Perfectly circular cells have a Circularity value of 1, with this decreasing as cells become more irregular/elongated. Filamentous cells, being elongated, have larger surface areas than normal planktonic *E. coli* cells. Their long cylindrical shape also means they are less circular than non-filamenting cells. We thus used these metrics to classify cells into filamentous and non-filamentous categories, and assessed their frequencies in RStudio (Posit Team, 2025). Unfocused cells were first filtered out. Using cells from the 0 CTX treatment 90-minutes post inoculation, we derived benchmark values for normal dividing cells. Those cells that were bigger than two standard deviations above the mean area of planktonic cells, as well as those that were two standard deviations below the mean circularity measure of planktonic cells were defined as filamentous. A random set of 400 cells were sampled for each concentration. For some higher CTX concentrations where this number of cells was not available, all imaged cells were sampled. Filament frequencies were then determined proportional to the total sampled cell population for each concentration at each timepoint.

### Model derivation

We model the dynamics of the number of CFUs over time semi-mechanistically as a function of the fraction of PBP3 and PBP1 that is not bound by antibiotic and is thus functionally active. The number of CFUs over time increases by cell division, and PBP3 is essential in that process. If PBP3 is fully inhibited, cell division does not occur, resulting in filamented cells that increase the biomass by growing in length, but not in cell number (CFU). Cell death is assumed to solely be due to failure to maintain the cell wall by PBP1, and hence the death rate increases as the concentration of unbound PBP1 decreases. This leads to the following equation:

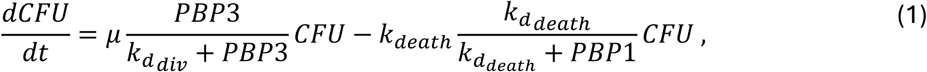

where μ is the maximum division rate, 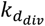 is the dissociation constant of PBP3 to the divisome, *k_death_* is the maximum death rate and 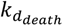 is the dissociation constant of PBP1 and the proteins/machinery required to maintain the cell. Both growth and death terms are modelled as Hill Equations with Hill-coefficient 1, as we assume there is no cooperative binding of PBP proteins to their substrates.

The fractions of unbound PBP1 and PBP3 depend on the periplasmic antibiotic concentration through a declining Hill equation, each with their own dissociation constant K_d_:

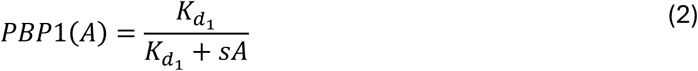

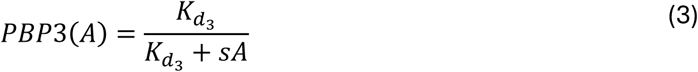

Here, s is a scaling factor which represents the ratio between external and periplasmic concentrations when modelling mutant dynamics. For the wildtype, we assume *s=1*, while for the mutant strains *s < 1* because these strains reduce periplasmic antibiotic concentrations by either reducing import, increasing export, or enzymatic degradation of antibiotic.

To reduce the number of parameters needed for fitting the model, we define 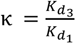 and 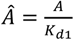, changing equations 2 and 3 into:

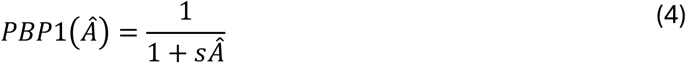

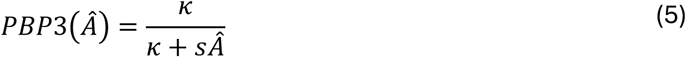

κ is the ratio of the affinities of both proteins towards antibiotic. Based on the reported affinities of cefotaxime to PBP1 and PBP3, we expect κ to be lower than 1.

### Parameter fitting

The model with 5 parameters: 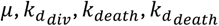 and κ, was fitted using the Optim.jl implementation of Simulated annealing (K Mogensen & N Riseth, 2018), which is a global optimization technique that is designed to avoid local optima by being able to accept worse solutions by some probability which lowers over time (Goffe, 1996; K Mogensen & N Riseth, 2018). Simulated annealing was performed with default parameters and a maximum of 1 million iterations. As starting parameters, 100 random sets were sampled using Latin hypercube sampling. Bounds for sampling and optimization were set to the values listed in Supplementary Table 1. κ was fixed according to the values found by Kocaoglu C Carlson (Kocaoglu C Carlson, 2015). Scoring of the model fit was done by calculating the variance weighted sum of squares of the data from both wild-type time-kill curves combined. The model from Regoes et al. was fitted using the same procedure, only with different bounds (Supplementary Table 2).

## Results

### Cefotaxime displays a two-step pharmacodynamic curve

To understand the relationship between the concentration of CTX and net rate of population change rates, time-kill assays were performed. Wild type (WT) bacteria were cultured over a range of CTX concentrations and sampled every hour for 3 hours to assess the change in CFU/mL. We observed a two-step decrease in net rate of change as CTX concentration increased, yielding a curve with three distinct phases (Fig. 1A). The first phase is defined by positive growth rates, spanning concentrations from 0 µg/mL up until 0.06 µg/mL. The average growth rate observed in this region was 0.29±0.127 hour^−1^. The second phase began at 0.12 µg/mL, as we approached the MIC (0.25 µg/mL, Fig. S2). Here, nearly no growth was observed (−0.09±0.06 hour^−1^), with CFU counts remaining almost constant over the 3 hours of the assay. In the third phase (concentrations of 7.8 µg/mL and higher), net rates become increasingly negative (<-0.2 hour^−1^) due to substantial killing of bacteria. The highest concentrations even showed complete bacterial clearance within the first 1-2 hours. The observed three-phase growth/kill dynamic differs from most data reported in previous work with β-lactams, where stable regions of growth and death were separated by a single “step” (Ferro et al., 2015; Foerster et al., 2015; Regoes et al., 2004). A second run of the assay, with a CTX concentration scale having 196 µg/mL as the maximum, showed very similar results (Fig. S1).

**Fig. 1:**
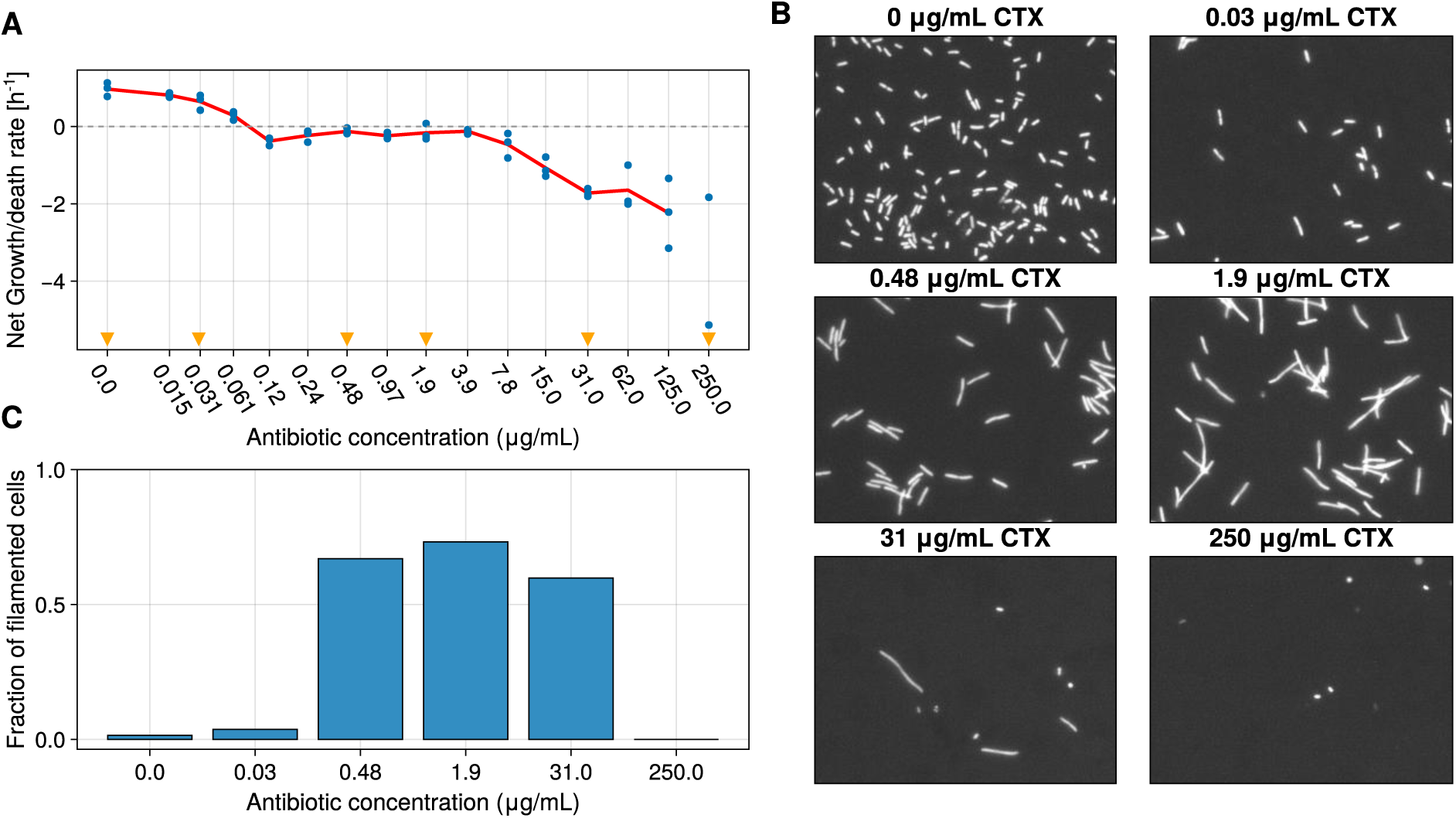
CTX shows a two-step pharmacodynamic curve against E. coli REL606. **(A)** Net rate of change of the bacterial population as a function of Cefotaxime (CTX) concentration. Points represent the independent biological replicates (n=3), with the mean plotted as a red line. Triangles on the x-axis indicate the concentrations at which cells were imaged using microscopy. **(B)** Representative microscopy images at each antibiotic concentration, after 90 minutes of incubation. C: The fraction of filamented cells at a subset of CTX concentrations spanning the three phases.

We suspected that the second phase (with approximately net 0 growth) resulted from filamentation, with cells continuing to grow, but not dividing under moderate CTX stress (Buijs et al., 2008; Gross et al., 2024). We tested this hypothesis using fluorescent microscopy (Fig. 1B). Changes in cell morphology were evaluated by culturing bacteria under 6 CTX concentrations spanning the 3 phases, sampling twice at 45-minute intervals and assessing morphologies via fluorescent microscopy. At both 45 and 90 minutes post inoculation, the 0 and 0.03 µg/mL CTX concentrations showed cylindrical dividing cells, supporting the positive growth rate of bacteria during the first phase (Fig. 1B). In the second phase, cells in 0.048 µg/mL CTX seemed unchanged at 45 minutes, while those in 1.9 µg/mL showed slightly elongated cells (Fig. S2). At 90 minutes post inoculation, both concentrations showed a population majorly comprised of filamented cells (Fig. 1B), with filaments making up 67% and 74% of the sampled cells at 0.048 µg/mL and 1.9 µg/mL CTX, respectively (Fig. 1C). In the third phase, at 31 µg/mL and 250 µg/mL CTX, we observed a population of smaller spherical cells, with some filaments at 31 µg/mL CTX (Figs. 1B & C). It should be noted that, at higher concentrations, substantial cell death meant fewer cells were available for imaging. We thus observe 3 distinct classes of morphology: healthy cells, filaments, and small spherical cells. Frequency changes in these associate with growth, maintenance, and death seen in the pharmacodynamic curve.

### A two-target binding model recapitulates the killing dynamics

To fit the pharmacodynamic models, we combined the data of the two time-kill assays, with different concentration ranges, that we performed for our wild-type strain. Fitting the model from Regoes et al. (Fig. 2B, grey line) resulted in a model that failed to capture the observed two-step function as expected (since this model assumes a single step in the decline of the net growth rate). As a result, this model underestimated the growth rate in the absence of antibiotic (0.37 hour^−1^ compared to 0.878±0.159 hour^−1^), and overestimated the MIC, which this model suggests is around 3.9 μg/ml while we experimentally determined it at 0.25 μg/ml (Fig. S2).

**Fig. 2:**
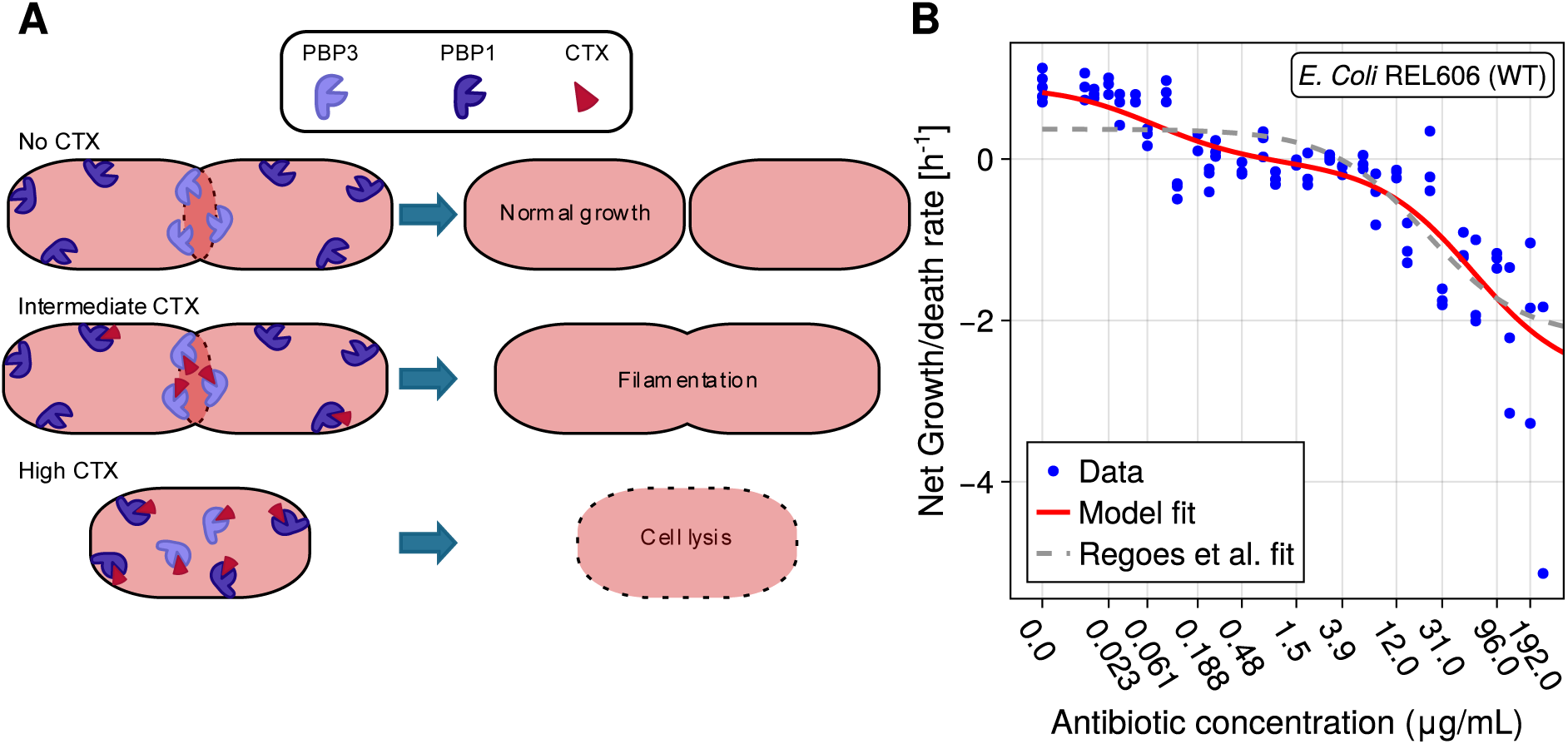
A semi-mechanistic model recapitulates the two-step growth/kill dynamics, improving fit. **(A)** Schematic of the putative mechanism behind the two-step dynamic observed in Fig. 1. At low antibiotic concentrations, cells can divide as normal. At intermediate concentrations, PBP3 (light purple), which has a high affinity for CTX (red triangles), is inhibited first. Inhibition of PBP3 prevents septation, resulting in filamented cells. PBP1 (dark purple), with lower affinity for CTX than PBP3, may be partially inhibited. At high CTX concentrations, PBP1 is also fully bound by antibiotic. Due to PBP1’s role in maintaining cell wall integrity, this causes lytic death. **(B)** Comparison of the fit of our two-step model and the single-step Hill function of Regoes et al. (2004) to the wild-type growth/kill data obtained from the two time-kill assays. Blue dots are the independent biological replicates (n=3), the grey dotted line is the predicted fit using the model from Regoes et al (Regoes et al., 2004), and the red solid line is the fit using our proposed semi-mechanistic model.

To explain the two-step killing rate function, we sought to make a semi-mechanistic model that incorporates the two targets of CTX: PBP3 and PBP1 (Fig. 2A), hypothesizing that the observed two-step behaviour is due to differences in the binding affinities and function of these two proteins. PBP3 has higher affinity for CTX and will be inhibited at lower antibiotic concentrations than PBP1 (Kocaoglu & Carlson, 2015). As PBP3 is bound, individual cells stop dividing and instead form filaments that grow in length (Hayes & Orr, 1983). Each filamented cell shows up as one colony forming unit in the experimental assay and likely does not burst in the time over which the assay was performed (Kjeldsen et al., 2015). This may explain the observed plateau with a net CFU growth rate around 0 in the second phase. Only at higher concentrations of CTX (phase 3), PBP1 also becomes inhibited, resulting in failure to maintain the cell membrane and subsequent lysis (Pazos & Vollmer, 2021; Yousif et al., 1985).

Based on these arguments, we created a model which consists of a cell division term dependent on PBP3, and a cell death term dependent on PBP1 (Equation 1), with the amount of free PBP proteins depending on the CTX concentration and their affinity for CTX (see Methods for full model derivation). Fitting this new model to the data, we now find a two-step decline in the growth rate with increasing antibiotic concentrations (Fig. 2B, red line). The growth rate in the absence of antibiotic is also estimated more accurately: the predicted growth rate is 0.826 hour^−1^ while the data show a growth rate of 0.878±0.159 hour^−1^.

### Resistance mutations alter the pharmacodynamic trend

Next to the wild-type strain, we also measured the time-kill kinetics of three mutants with varying resistance mechanisms (Fig. 3A): (1) *ΔompF:* a knock-out of a porin, reducing the permeability of CTX; (2) *ΔacrR*: a knock-out of a repressor of the efflux pump AcrAB-TolC, resulting in the overexpression of these efflux pumps; and (3) a mutant expressing plasmid-encoded TEM1-G238S, a β-lactamase enzyme with high affinity for CTX over wild type TEM1.

**Fig. 3:**
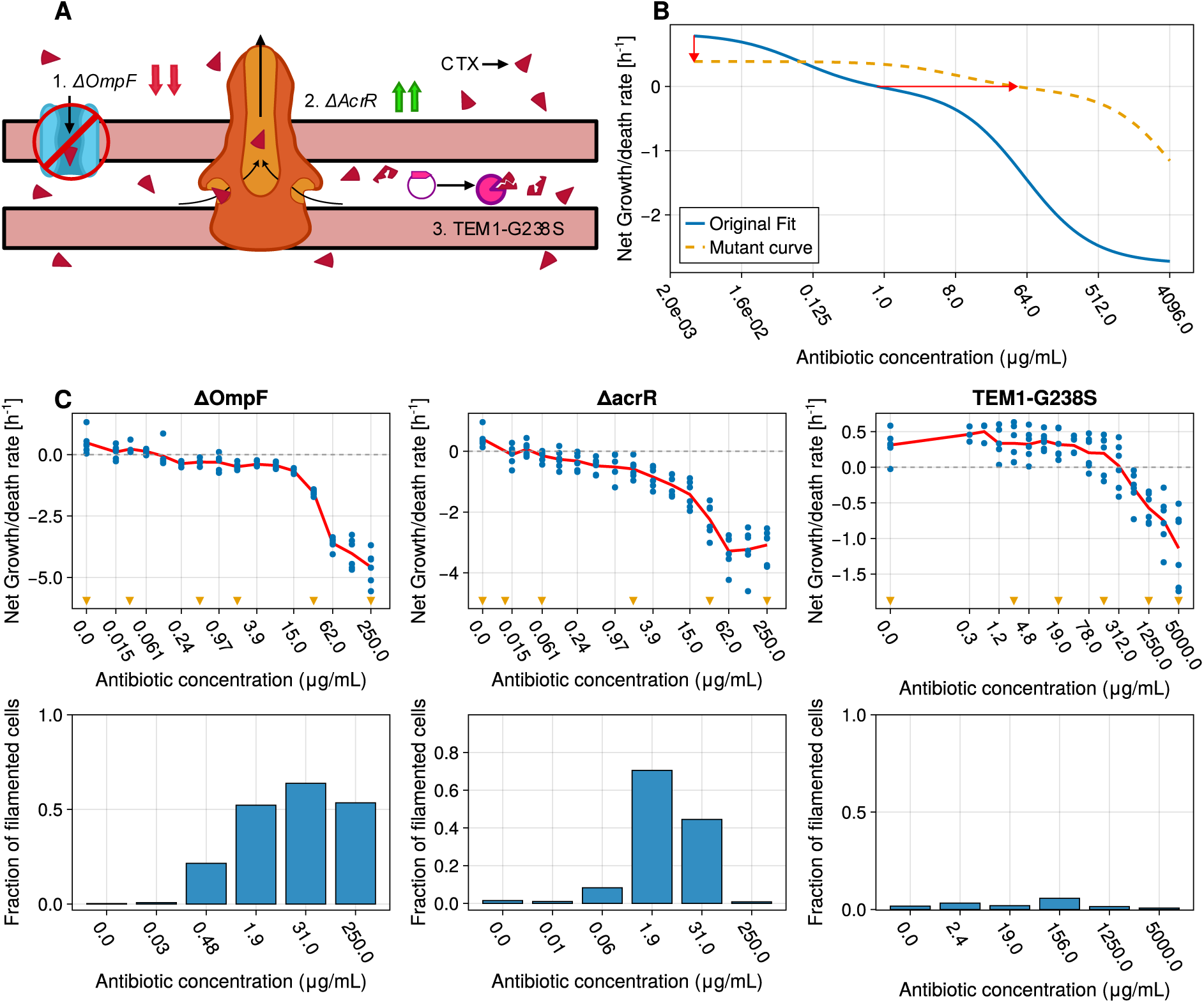
Growth/kill dynamics of resistant mutants are different from the WT and vary with each mutant strain. **(A)** Schematic illustrating the mutant strains for which time-kill kinetics were evaluated: 1. ΔompF, which lacks the ompF porin, resulting in a reduced permeability of CTX. 2. ΔacrR, which has transcription factor acrR deleted, which results in upregulation of the AcrAB-TolC efflux pump. 3. A mutant expressing TEM1-G238S beta-lactamase, enabling the degradation of CTX. **(B)** Model prediction in a possible mutant strain, together with wild-type as blue line. The yellow dashed line shows a predicted time-kill curve for a mutant with a fitness cost and reduction of periplasmic antibiotic concentration (μ = 0.5, s=0.01). The red arrows indicate the drop in growth rate in absence of antibiotic and the shift in antibiotic concentration at which the net growth rate is zero. **(C)** rate of change of mutant population as a function of antibiotic concentration (top), together with estimated filamentation fractions from microscopy images (bottom). Blue dots are independent replicates (n=6), dark red line represents the mean.

The resistant mutants all influence the intracellular (periplasmic) concentration of CTX. To get an idea of what the pharmacodynamic curve of the mutants may look like, we adapted the model and added a scaling parameter *s* that can be used to modulate the difference between extracellular CTX and periplasmic CTX. We then simulated a pharmacodynamic curve of an arbitrary mutant, with a reduced growth rate and a reduction in periplasmic CTX (Fig. 3B). This resulted in a curve where the two-step behaviour was less pronounced, where the reduction in growth rate lowers the first phase, and the reduction in periplasmic CTX shifts the point where the growth rate becomes zero to higher concentrations as expected. However, the transition phase is also more drawn out.

Time-kill assays and morphological assessments of the mutant strains were conducted in the same fashion as the wild-type. All 3 mutant strains incurred fitness costs, showing lower growth rates than the WT (0.42 ± 0.07 hour^−1^) when CTX was absent (*ΔompF* = 0.2 ± 0.19 hour^−1^, *ΔacrR* = 0.17 ± 0.12 hour^−1^, G238S = 0.13 ± 0.09 hour^−1^). The *ΔompF* mutant showed similar dynamics to the WT: a two-step function with phases of growth, stasis, and death (Fig. 3C). Deviation from the WT was observed in the first 2 phases: the growth cost meant lower growth rates than the WT over the first phase (0.1±0.13 hour^−1^), causing the first step to be less pronounced here, while growth rates in the second phase were slightly negative (−0.17±0.08 hour^−1^). The deletion of the OmpF porin also seems to have lessened the extent of filamentation in this strain. Where the WT showed high filament frequency (67 %) at 0.48 µg/mL CTX, we saw little filamentation in *ΔompF* at this concentration (19%) (Fig. 3C). Further, the highest degree of filamentation was observed at 31 µg/mL CTX (64 %) as compared to 1.9 µg/mL CTX in the WT.

*ΔacrR*, the export mutant, deviated completely from the two-step function. While the no drug control did show a positive growth rate of 0.17±0.13 hour^−1^, almost none of the cultures containing CTX showed any growth (Fig. 3C). Growth rates gradually declined with each doubling in CTX concentration until 15 µg/mL after which the decline was sharper. *ΔacrR* did not markedly differ from the WT in terms of filamentation behaviour (Fig. S4), with both showing highest filamentation at 1.9 µg/mL.

The G238S strain behaved differently from any of the other strains (Fig. 3C). To account for the greater resistance of this strain, CTX concentrations up to 5000 µg/mL, rather than 250 µg/mL were used. This strain, strikingly, showed positive growth rates until 312 µg/mL; following which the net rate of change decreased until 5,000 µg/mL. No filamentation was observed in the G238S strain, in the assessed concentrations, likely due to the strong effect of the mutated TEM-1 β-lactamase on CTX resistance (Fig. S6).

We next asked whether the resistant mutants’ pharmacodynamic curves could be predicted from the wild-type model alone. To the wild-type parameters, we applied two adjustments: a reduction in maximum growth rate and the addition of a parameter defining the intracellular concentration of antibiotic, reflective of the resistance mutation. Given the way the model is set up (see supplementary information for more details), a correction had to be made to allow for these adjustments. Further, since the MICs of *ΔompF and ΔacrR are close to that of the WT, the additional parameter for reduced intracellular concentration was only of significance for G238S.* The resulting curves did not fit the experimental data well (Fig. 4). For *ΔompF*, the model recapitulates the data well from 0 μg/mL up to 15 μg/mL. At higher concentrations, however, the model underestimates the death rate and the predicted curve has a lower slope than the data. For the *ΔacrR* mutant, the model consistently overestimates the net rate of change, with the discrepancy widening at higher concentrations of CTX. The G238S strain, like Δ*ompF* fits the data at lower concentrations well but not at higher concentrations. Together, these deviations suggest that resistance alters the shape of the concentration–response relationship rather than simply repositioning the wild-type curve along the concentration and growth-rate axes. We also performed a free fit of the model, without constraining the parameters to their wild-type values, to test whether an alternative combination could better explain the data. This yielded a higher estimated maximum death rate than in the wild type. Although the maximum growth rate was estimated at 1 for all mutants (the cap in the model), the growth cost of resistance was still captured by the model through the other fitted parameters (see Supplementary Information).

**Fig. 4:**
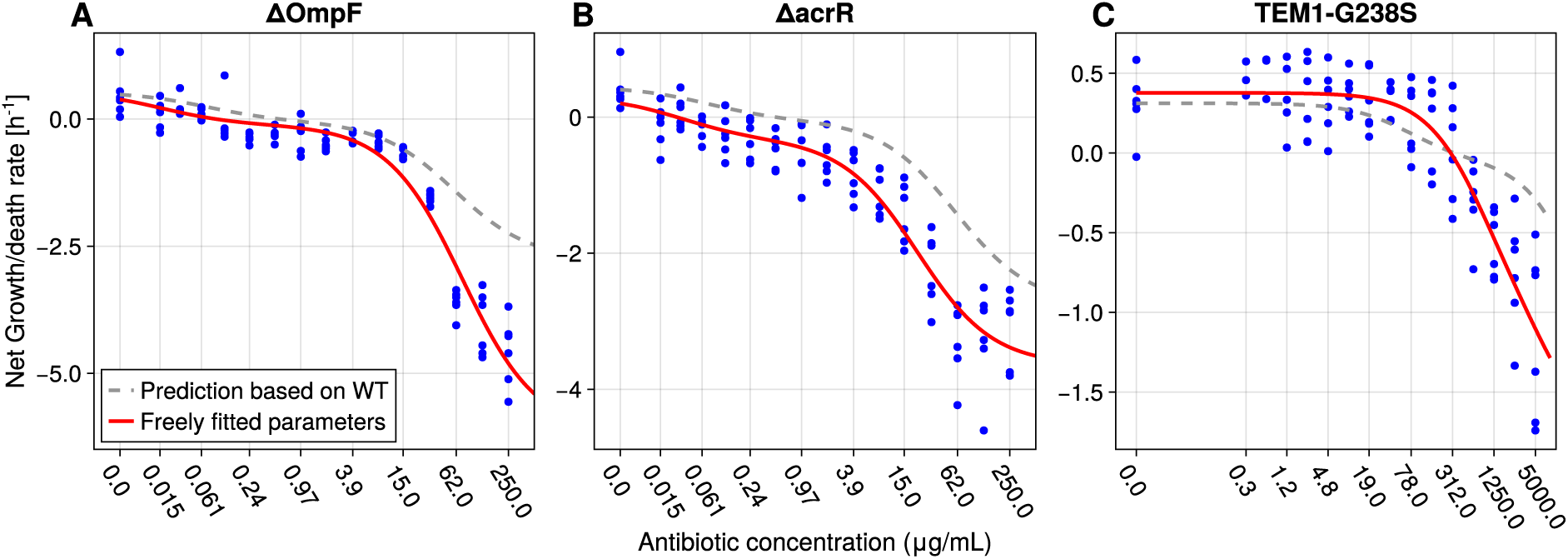
Growth costs and resistance mechanism do not fully explain altered pharmacodynamic trend of resistant mutants. Predicted pharmacodynamic curves for each resistant strain using the WT model with adjusted parameters are plotted as a grey dashed line., The model with freely fitted parameters is represented by the red line. Blue dots are independent experimental replicates (n = 6).

## Discussion

In this study, we aimed to understand killing by the β-lactam antibiotic cefotaxime (CTX) by characterizing its pharmacodynamic curve against *E. coli* REL606 and derived CTX resistant mutants. Contrary to the classically reported sigmoidal response of the net rate of change with antibiotic concentration (Regoes et al., 2004), we found that the wild-type killing of *E. coli* REL606 by CTX follows a two-step pharmacodynamic curve. A semi-mechanistic model incorporating the interaction between CTX and its target penicillin binding proteins (PBP) was able to explain the observed data. Binding of CTX to its primary target PBP3 induced filamentation that conferred antibiotic tolerance stopping further reduction in population rate of change. Only at higher concentrations of antibiotic, when PBP1 was bound, did the cells start lysing resulting in the second step.

Similar two-step behaviour has been reported previously. Foerster et al. in 2016 found that Chloramphenicol, Ceftriaxone, Cefixime, and Benzylpenicillin had similar two-step curves against *Neisseria gonorrhoeae* (Foerster et al., 2016). While they did acknowledge the two-step behaviour, noting that it might indicate distinct targets, they excluded the second step from their analyses in order to use the classical pharmacodynamic function to fit their data. All three β-lactams in their dataset bind primarily to PBP2 at lower concentrations followed by binding to PBP1 at higher concentrations (Barbour, 1981). PBP2 and PBP1 in *N. gonorrhoeae* are homologues of PBP3 and PBP1 in *E. coli* (López-Argüello et al., 2023) with demonstrated function for PBP2 in septum formation (Zou et al., 2017). Although, *N. gonorrhoeae* is a diplococcus and cannot filament, inhibition of PBP2 does cause the cells to swell up over time (Westling-Häggström et al., 1977). Given the target homology, shared function and the correspondingly similar two-step pharmacodynamic curves, their results are consistent with the mechanism we describe here.

The mutations studied are the most frequent to evolve in response to CTX (Schenk et al., 2022). We aimed to understand how they would affect the pharmacodynamic curve observed in the WT. *ΔompF* showed a pharmacodynamic function most like the WT, though the first step was less pronounced and growth rates in the second phase were lower than the WT. This indicates a cost of resistance. *ΔompF, further*, showed far lower filamentation at 0.48 µg/mL as compared to the WT, and surviving filaments were imaged at higher concentrations (31 µg/mL and 250 µg/mL) where the WT showed almost complete clearance. This indicates towards the OmpF porin’s deletion likely playing a role in delaying the binding of CTX to PBPs by reducing antibiotic influx. *ΔacrR* differed from all the other strains in its pharmacodynamic curve. Except for the no drug control, all other tested concentrations had a net rate of population change of 0 or lower. Mutational costs do completely not explain this behaviour as the no drug control showed net positive growth although lower than that of the WT. As *acrR* encodes a transcriptional regulator, its deletion may exert pleiotropic effects beyond de-repression of the AcrAB efflux pump, which could contribute to the altered pharmacodynamic response.

The G238S strain did not display a two-step pharmacodynamic curve in the tested concentration range. Fittingly, no filamentation was observed either. Owing to the very large concentrations of CTX tested in 2-fold intervals, we may have missed the concentrations where G238S would filament. The concentrations assessed thus likely capture the growth phase where neither PBP system is inhibited, following which a single doubling of concentrations would bring us to the third phase, where both PBP3 and PBP1 systems are bound and cell lysis occurs. Time-kill assays with increased resolution between 19 µg/mL and 312 µg/mL would be required to test this.

Model parameters from the fitted WT data when used with adjustments for the resistance mechanism, could not fit the mutant data well. This is likely due to our inability to accurately measure the maximal death rate, *k_death_ in vitro* for the WT. This could be caused by two factors. First, the large number of concentrations assessed restricted sampling to once every hour, limiting the temporal resolution of the assay. Second, at higher concentrations, there was substantial carry-over of CTX in the plated cultures, clearing bacteria post-sampling on the plate. As a result, lower dilutions could not be measured, effectively reducing our limit of detection 100-fold. Owing to their resistance to CTX, the effect of on-plate killing was likely dampened in the mutant strains, improving detection at smaller CFU counts relative to the WT. Given the additional resolution we could better estimate lower CFU values of the mutant giving rise to a steeper slope than for the WT. The value of *k_death_* derived from fitting the WT data thus overestimated the mutant maximum death rate. Assessing fewer concentration with greater temporal resolution or adding an efficient β-lactamase to the spotting plate, mitigating on-plate killing, could help us estimate *k_death_* more accurately.

β-lactams are widely considered to be time-dependent, with efficacy tied to the duration antibiotic concentrations stay above the MIC (Brogden & Spencer, 1997; Gunderson et al., 2001; Vogelman et al., 1988). Our observations, however, show a partly concentration-dependent effect in the case of CTX acting against *E. coli*. In what we define as the second step, at concentrations greater than 3.9 µg/ml, the rate of bacterial killing increases with each doubling of CTX as increasing amounts of PBP1 is bound. It is to be noted, though, that part of the data would also be consistent with a time dependent kinetic. The filaments would eventually burst after reaching a critical length which would happen with longer time under antibiotic (Hayes & Orr, 1983; Kim et al., 2023; Tuomanen et al., 1986). There are multiple reasons existing pharmacodynamic studies may miss a two-step dynamic. Most studies do not test an extended range of antibiotic concentrations like we did (Ronaghinia et al., 2026) and if they do, the data is analysed using linear regression (Bonapace et al., 2002; Lutsar et al., 1997; Vogelman et al., 1988) – a framework incapable of detecting non-linear two-step behaviour. Indeed, a study that analysed data split before and after a concentration threshold found differences in bactericidal activity that became marked after a threshold (Erlendsdottir et al., 2001). The two-step behaviour that we observe for CTX compels us to re-evaluate how its antibacterial activity is understood.

The existence of an intermediate phase where bacteria tolerate the antibiotic due to filamentation has implications for resistance evolution. Bacterial filaments still replicate their chromosomal DNA even if septation is stalled due to β-lactam exposure (Cayron et al., 2023). This could allow for the generation of mutations that may confer resistance. If the antibiotic treatment elicits an SOS response, mutation rates may increase by several orders of magnitude increasing the likelihood of a resistance mutation evolving (Bos et al., 2015; Cirz et al., 2005). This would be consequential for intermittent antibiotic infusion therapies. If, between doses, the concentration were to fall below a threshold, filaments could resume division (Cayron et al., 2023; Zahir et al., 2020), allowing the acquired resistance mutations to proliferate as new lineages and be selected for. Even if resistance evolution did not occur, the reduction in antibiotic concentration could still recover a part of the original population. Serum concentrations targeted during antibiotic treatment are usually between 1-8 times the MIC (Aardema et al., 2020; Guilhaumou et al., 2019). In the case of *E. coli* REL606 and CTX, our data shows that this still lies in the region of filamentation, posing the potential risk of expediting AMR evolution. Consideration of such complex dynamics in evaluating therapeutic concentrations used could prove important in combating the threat of AMR evolution in the long-term.

Understanding how bacteria respond to varying antibiotic concentrations is essential for designing effective dosing strategies that maximize treatment success and limit resistance evolution. Here, we report an unusual two-step pharmacodynamic curve for the β-lactam cefotaxime, and extend the classical pharmacodynamic model by incorporating a mechanistic layer that explains our data. Future work could take a mechanism-first approach, testing whether antibiotics with a target profile similar to cefotaxime’s exhibit comparable pharmacodynamics, and, conversely, whether β-lactams that do not share this profile follow more canonical dynamics. Such comparisons would help to solidify the link between an antibiotic’s mechanism of action and its pharmacodynamic profile.

## Acknowledgements

We would like to thank Marloes Meeus and Francisca Reyes Marquez for technical help, Vaughn Cooper for providing CFP labelled *E. coli* REL606, and Philip Ruelens and Clement Bijl for constructing the resistant mutants in that background.

## Funding

This work was funded by the Netherlands Organisation for Scientific Research grant OCENW.M.22.248 and the Collaborative Research Center 1310 of the German Research Foundation.

## Author contributions

J.P.: Visualization, Formal Analysis, Writing – original draft, Writing - review & editing, Conceptualization, Investigation, Methodology

A.R.: Formal Analysis, Writing – original draft, Writing - review & editing, Investigation, Methodology

J.A.G.M: Funding acquisition, Supervision, Writing - review & editing, Conceptualization H.M.D.: Funding acquisition, Supervision, Writing - review & editing, Conceptualization

A. B.: Supervision, Writing – review & editing, Conceptualization, Investigation, Methodology

## Data availability

The data that were used for the findings of the study, including code for the model and figures, can be found at https://git.wur.nl/pool024/pbpmodel.

## Notes

https://git.wur.nl/pool024/pbpmodel

### Competing Interest Statement

The authors have declared no competing interest.

