## Supplementary Information for "Antibiotic tolerance due to filamentation shapes β-lactam pharmacodynamics in *Escherichia coli*"

### μ and its relationship with the growth rate in absence of antibiotic

By modelling the unbound proteins as a fraction, the input for the Hill equations that the proteins are variables of, lies between 0 and 1. Usually it is the case that the limit goes to infinity. This results in a discrepancy between model parameter μ, the maximum growth rate, and the actual growth rate after evaluating the model at zero antibiotic. At zero antibiotic, both PBP3 and PBP1 fractions are one. When we fill this in, we get equation 6 below, which gives us the relation between the net growth rate at 0 and μ.

$$\mu_{A_0} = \mu \frac{1}{k_{d_{div}}} - k_{death} \frac{k_{d_{death}}}{k_{d_{death}} + 1} \quad (1)$$

This equation can be rewritten to express μ in terms of $\mu_{A_0}$:

$$\mu = \frac{\mu_{A_0} + k_{death} \frac{k_{d_{death}}}{k_{d_{death}+1}}}{\frac{1}{k_{d_{div}}}} \quad (2)$$

We can use this equation to ensure the growth rate at zero antibiotic matches with the value obtained from the data. We used this correction in predicting the time-kill curves for the mutants.

With the fitted parameters, the resulting μ for the wild-type and mutants is around 5-15% higher than the mean growth rate of the data. This 5-15% difference is well within the standard deviation of the actual data.

*Supplementary Table 1: Parameters used in the two-target model, with their biological description, unit, and their value resulting from fitting the WT data. The upper and lower bounds used in Latin hypercube sampling and Simulated annealing are also given.*

| Parameter | Description | Unit | Value | Lower bound | Upper bound |
|---|---|---|---|---|---|
| $\mu$ | Maximum growth rate | $hour^{-1}$ | 1.0 | 0 | 1.0 |
| $k_{death}$ | Maximum death rate | $hour^{-1}$ | 2.7375639 | 0 | 100 |
| $\kappa$ | Affinity ratio of PBP proteins to antibiotic | 1 | 0.0001242 | Fixed | Fixed |

| | | | | | |
|---|---|---|---|---|---|
| $k_{d_{div}}$ | Affinity of PBP3 to the divisome | 1 | 0.00203911 | $1*10^{-5}$ | 1000 |
| $k_{d_{death}}$ | Affinity of PBP1 to 'maintenance' reaction | 1 | 0.0176961 | $1*10^{-5}$ | 1000 |

*Supplementary Table 2: Bounds set for optimizing the two-target model using Latin hypercube sampling and Simulated annealing*

| Parameter | Lower bound | Upper bound |
|---|---|---|
| $\mu$ | 0 | 1.0 |
| $k_{death}$ | 0 | 100 |
| $\kappa$ | Fixed | Fixed |
| $k_{d_{div}}$ | $1*10^{-5}$ | 1000 |
| $k_{d_{death}}$ | $1*10^{-5}$ | 1000 |

*Supplementary Table 3: Parameters used in the model from Regoes et al., with their biological description, unit, and their value resulting from fitting the WT data. The upper and lower bounds used in Latin hypercube sampling and Simulated annealing are also given.*

| Parameter | Description | Unit | Value | Lower bound | Upper bound |
|---|---|---|---|---|---|
| $\psi_{max}$ | Maximum net growth rate | $hour^{-1}$ | 0.3733 | 0 | 1.0 |
| $\psi_{min}$ | Minimal net growth/death rate | $hour^{-1}$ | -2.1997 | -100 | -100 |
| $zMIC$ | Pharmacodynamic MIC | $\mu g * mL^{-1}$ | 3.657 | 1*10^-5 | 1000 |
| $\kappa$ | Hill coefficient | 1 | 1 | 1 | 20 |

*Supplementary Table 4: Bounds set for optimizing the model from Regoes et al. using Latin hypercube sampling and Simulated annealing*

| Parameter | Lower bound | Upper bound |
|---|---|---|
| $\psi_{max}$ | 0 | 1.0 |
| $\psi_{min}$ | -100 | -100 |
| $zMIC$ | $1*10^{-5}$ | 1000 |
| $\kappa$ | 1 | 20 |

*Table 1: Parameters obtained after optimization of the model for the data of each strain respectively. Reported values are the mean and standard deviation of the parameter sets in the best optimum, rounded to 4 digits. The standard deviation was omitted when rounded to 0. $\mu_0$ was calculated from the obtained parameters (see Supplementary)*

| Strain | $\mu_0$ | $\mu$ | $k_{death}$ | $k_{d_{death}}$ | $\kappa$ | $k_{d_{div}}$ | $s$ |
|---|---|---|---|---|---|---|---|
| Wild type | 0.825 | 1.0 | 2.7333 | 0.0178 | 0.0111 | 0.1458 | 1 |
| *ΔompF* | 0.3874 | 1.0 | 6.1784 | 0.0192 | 0.0111 | 0.9856 | 0.7261 |
| *ΔacrR* | 0.2033 | 1.0 | 3.6459 | 0.0996 | 0.0111 | 0.8739 | 0.515 |
| G238S | 0.3791± 0.0012 | 0.8137± 0.2508 | 1.8432± 0.1759 | 0.3053±0.18 | 0.0111 | 0.0152±0.0165 | 0.0247 |

*Table 2: Best parameter set obtained by fitting the data for each strain*

| Strain | $\mu_0$ | $\mu$ | $k_{death}$ | $k_{d_{death}}$ | $\kappa$ | $k_{d_{div}}$ | $s$ |
|---|---|---|---|---|---|---|---|
| Wild type | 0.8255 | 1.0 | 2.7624 | 0.0172 | 0.0111 | 0.1466 | 1 |
| *ΔompF* | 0.3873 | 1.0 | 6.1784 | 0.0996 | 0.0111 | 0.9856 | 0.7260 |
| *ΔacrR* | 0.2033 | 1.0 | 3.6459 | 0.0191 | 0.0111 | 0.8739 | 0.515 |

| | | | | | | | |
|---|---|---|---|---|---|---|---|
| G238S | 0.3775 | 1.0 | 1.9274 | 0.4767 | 0.0111 | $3.224*10^{-4}$ | $3.822*10^{-3}$ |

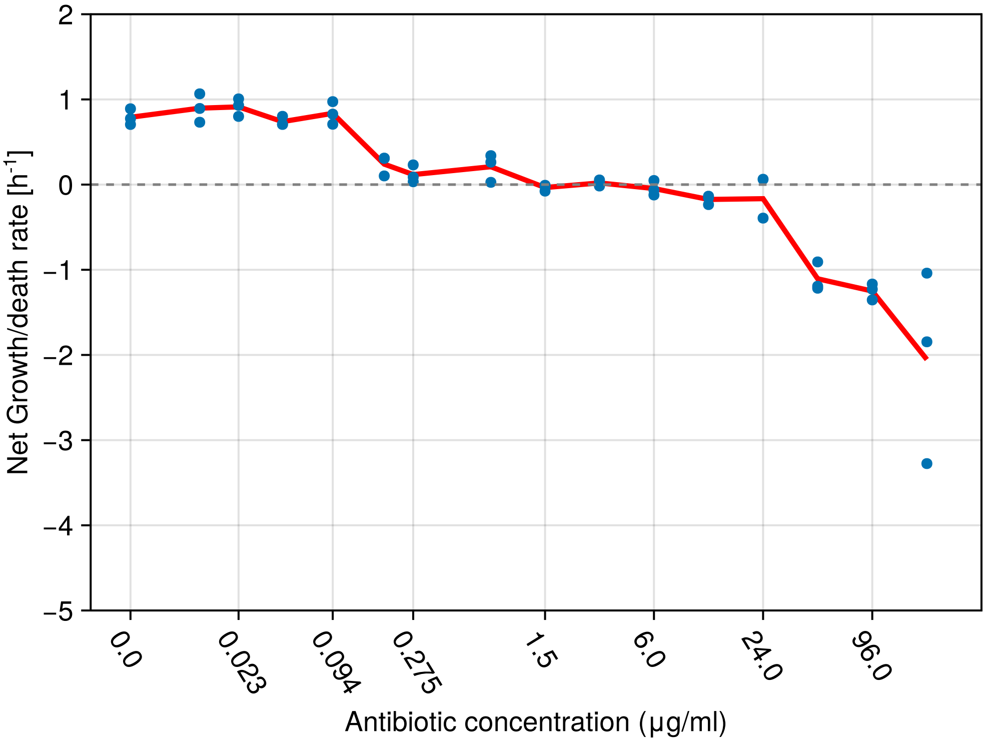


*Supplementary Figure 1: Another time-kill curve assay for the wild-type strain, which shows the same observed two-step dynamics.*

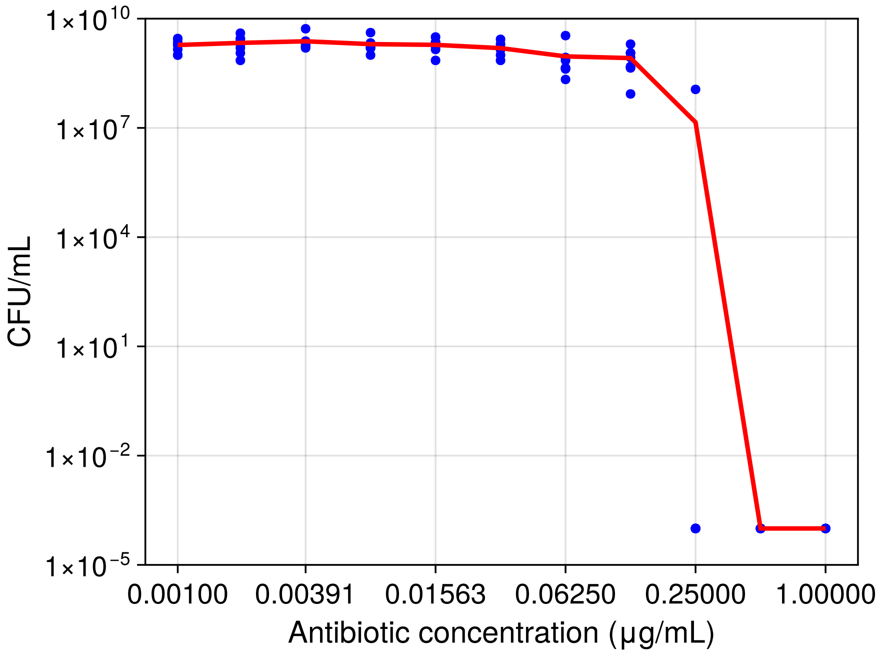


*Supplementary Figure 2: CFU based broth microdilution assay performed for the WT strain. 10 concentrations were tested up to 1 µg/ml, including a condition without antibiotic (here pictured at 0.001 µg/ml due to log-axis limitations). $8*10^5$ cells were incubated for 22 hours. CFU was determined by spotting dilution assay. Plotted is the mean of 6*

*replicates (3 biological x 2 technical) and individual data points. Three replicates at 1 μg/ml were removed due to showing growth, likely due to contamination. MIC was determined around 0.25μg/ml, as 5/6 replicates showed no growth.*

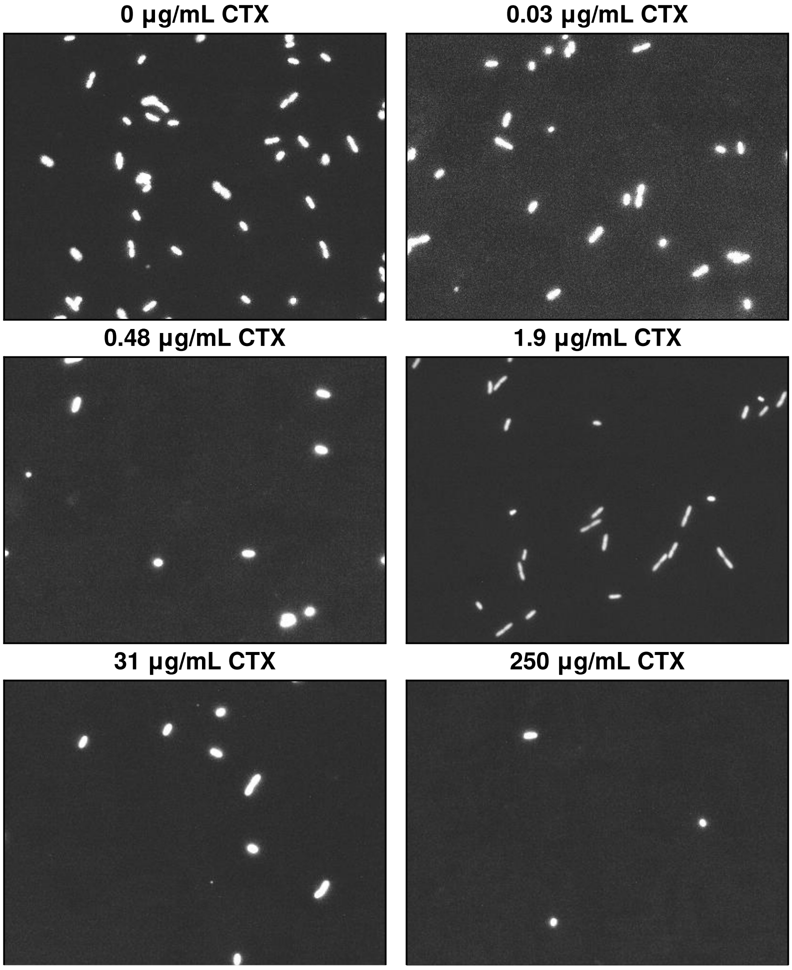


*Supplementary Figure 3: Representative microscopy images of the wild-type strain after 45 minutes of incubation in their respective CTX concentration.*

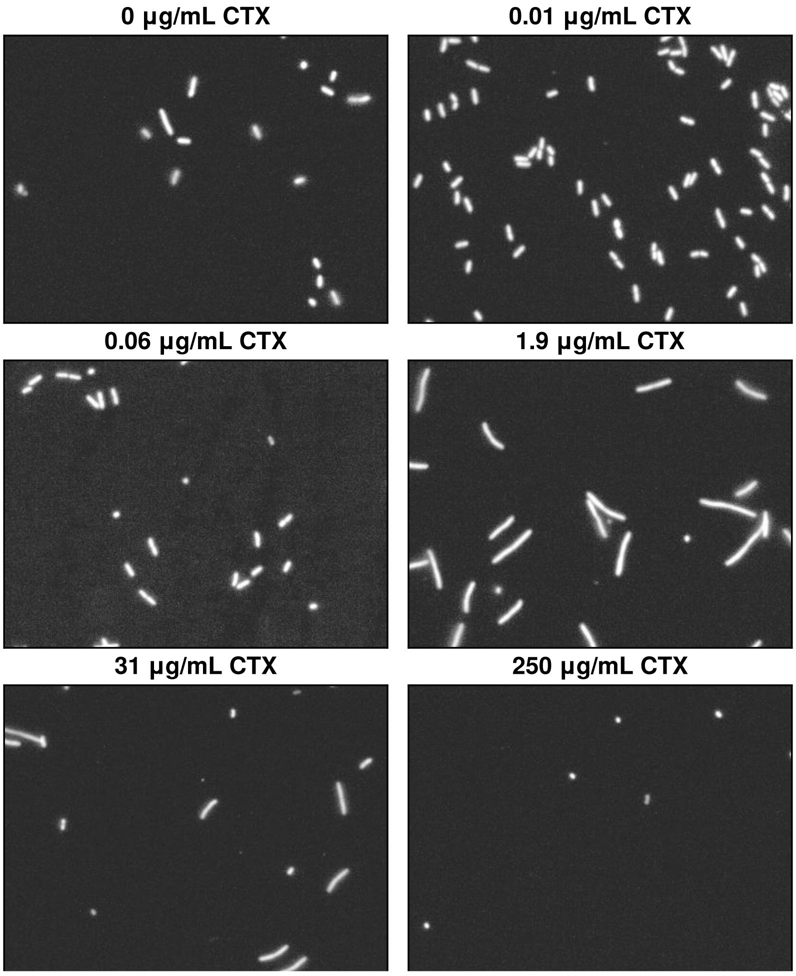


*Supplementary Figure 4: Representative microscopy images of the ΔacrR strain after 90 minutes of incubation in their respectivkjele CTX concentration.*

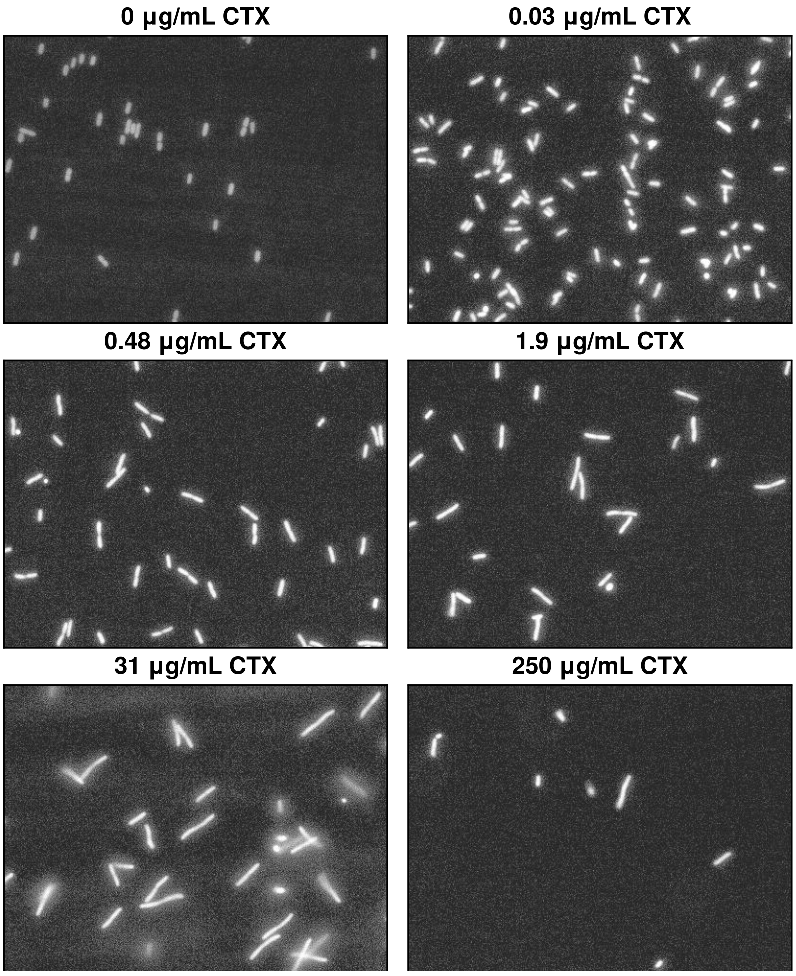


*Supplementary Figure 5: Representative microscopy images of the ΔompF strain after 90 minutes of incubation in their respective CTX concentration.*

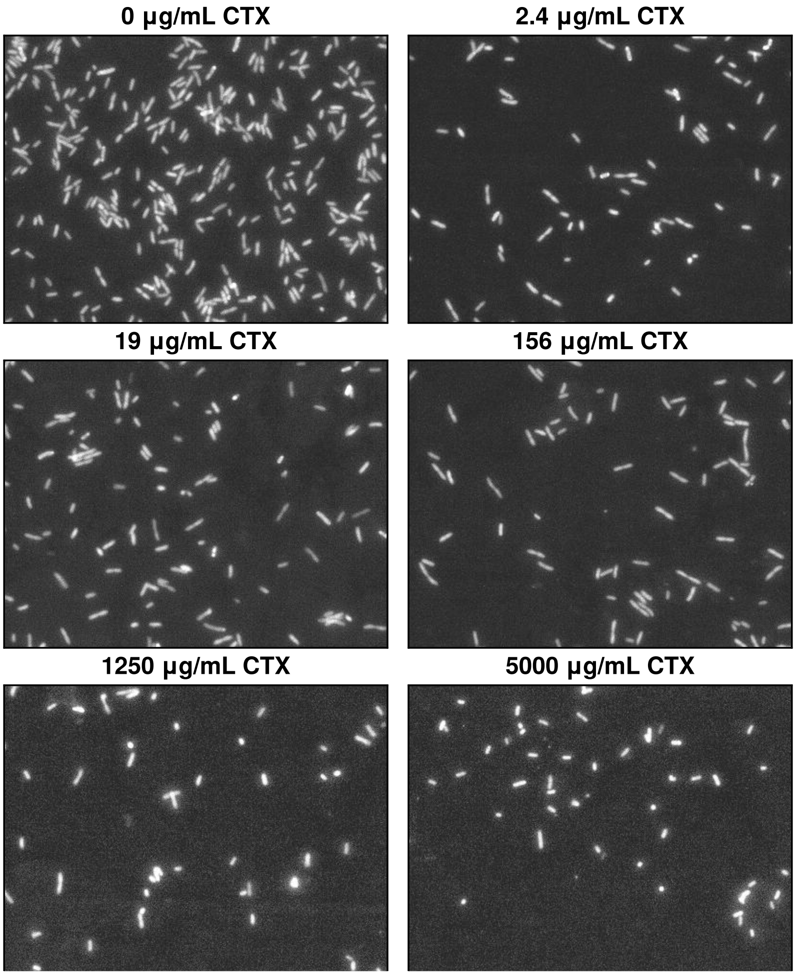


*Supplementary Figure 6: Representative microscopy images of the TEM1*-G238S *strain after 90 minutes of incubation in their respective CTX concentration.*
